# VarMatch: robust matching of small variant datasets using flexible scoring schemes

**DOI:** 10.1101/062943

**Authors:** Chen Sun, Paul Medvedev

## Abstract

**Motivation:** Small variant calling is an important component of many analyses, and, in many instances, it is important to determine the set of variants which appear in multiple callsets. Variant matching is complicated by variants that have multiple equivalent representations. Normalization and decomposition algorithms have been proposed, but are not robust to different representation of complex variants. Variant matching is also usually done to maximize the number of matches, as opposed to other optimization criteria.

**Results:** We present the VarMatch algorithm for the variant matching problem. Our algorithm is based on a theoretical result which allows us to partition the input into smaller subproblems without sacrificing accuracy VarMatch is robust to different representation of complex variants and is particularly effective in low complexity regions or those dense in variants. VarMatch is able to detect more matches than either the normalization or decomposition algorithms on tested datasets. It also implements different optimization criteria, such as edit distance, that can improve robustness to different variant representations. Finally the VarMatch software provides summary statistics, annotations, and visualizations that are useful for understanding callers’ performance.

**Availability:** VarMatch is freely available at: https://github.com/medvedevgroup/varmatch

## 1 INTRODUCTION

In recent years, next-generation sequencing data has been used in medical and genetic research to identify how genome mutations are related to phenotypes of interest (1000 Genomes Project Consortium *et al.*, 2012). In most of the studies, small variant calling, including the detection of single nucleotide variants (SNVs), multiple nucleotide variants (several SNVs occuring next to each other), or small indels (usually less than 30bp), plays a significant role. Small variant calling is a mature area, with several state-of-the-art tools, such as FreeBayes (Garrison and Marth, 2012), GATK (McKenna *et al.*, 2010), SAMtools (Li *et al.*, 2009), SNVer (Wei *et al.*, 2011), Platypus (Rimmer *et al.*, 2014), VarScan (Koboldt *et al.*, 2009), and Isaac (Raczy *et al.*, 2013). Detected variants are represented using the VCF file format (Danecek *et al.*, 2011).

An important starting point of many downstream analyses is to compare two VCF files to each other, to find matching variants. This is important for 1) measuring the similarity and population structure of several genomes (1000 Genomes Project Consortium *et al.*, 2010), 2) checking that the new variants added to a database do not already exist there (Assmus *et al.*, 2013; Tan *et al.*, 2015), 3) generating a high-confidence variant set by taking the intersection of the results of different variant callers (Zook *et al.*, 2014), and 4) evaluating the relative accuracy of different tools (Baes *et al.*, 2014) and understanding the source of their errors (Li, 2014). There have been several studies comparing datasets on the same genome generated by different aligners and variant callers (Cheng *et al.*, 2014; Baes *et al.*, 2014; Li, 2014; Hwang *et al.*, 2015; Cornish and Guda, 2015; Highnam *et al.*, 2015), and there is various software available to identify matching variants in two VCF files (`vcftools, rtgtools, bcftools, vt, bcbio, SMaSH` (Talwalkar *et al.*, 2014)).

Unfortunately, identifying matching variants in two VCF files is not as simple as may first seem, because applying two different VCF entries to a genome may result in the exact same donor sequence (they are *equivalent*). A VCF entry gives an allele sequence, its position on the reference, one or more alternate allele sequences of the donor, and, possibly, the donor genotype. The straightforward *strict matching algorithm* matches VCF entries which are identical, i.e. two entries that have the same position and the same reference and alternate alleles. However, this algorithm fails to match equivalent entries which are not identical. For example, Figure 1(a) illustrates how the same 2 bp deletion can be represented by four different VCF entries.

**Fig. 1:**
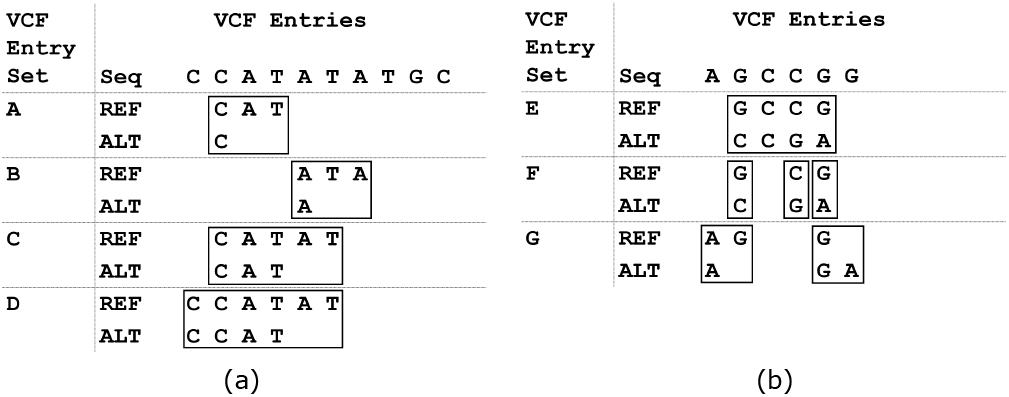
Examples of how the same variants can be represented by different VCF entries. VCF entries are represented by boxes but are grouped together into entry sets A-G. Panel (a) illustrates a variant which is a deletion of an AT from a short tandem repeat. VCF entry sets A-D are all singletons which are equivalent and represent this deletion. Panel (b) illustrates a complex variant which replaces the reference sequence GCCG with CCGA in the donor. This variant is represented by three equivalent but non-identical VCF entry sets (E-G). E is a singleton, F is composed of three entries, and G is composed of two entries. F is the normalized decomposition of E, and G is already normalized and cannot be decomposed further. The normalization algorithm would not detect any match, while the decomposition algorithm would not match G to E or F.

One way to address this problem is *normalization*. Tan *et al.* (2015) showed that there is a canonical way to represent VCF entries such that two entries are equivalent if and only if their canonical representations are identical. The *normalization algorithm* is to first normalize every entry and then run the strict matching algorithm. Normalization guarantees to identify equivalent pairs of VCF entries. However, a single variant can be represented by different non-singleton sets of VCF entries. Figure 1(b) illustrates three possible ways to represent the same variant. Each individual entry is normalized, but the three VCF entry sets are not identical even though they are equivalent. These entries will not be matched by the normalization algorithm. Such variants are called *complex*.

The `vcflib` package provides a way to partially address this problem through the decomposition of complex variants. It uses alignment of alternate allele to reference allele to break-up, or decompose, a complex VCF entry into multiple shorter ones. The *decomposition algorithm* proposed in (Li, 2014; Zook *et al.*, 2014) is to first decompose all entries, then normalize them, and then run the strict matching algorithm. Decomposition can help match some VCF entry sets, but it still does not work in some cases (see example in Figure 1(b)) Moreover, the decomposition varies based on the alignment, which is sometimes not unique or is not provided (e.g. in Platypus). Decomposition also allows fractional matches of a variant, which is difficult to interpret biologically For example, a complex variant can be decomposed into three smaller variants of which only one is matched.

An alternate approach, which we also take in this paper, avoids strict matching altogether. To check if two sets of entries are equivalent, we just apply them to the reference and check if the resulting donor sequence is the same. Matching two variant datasets can then be formulated as finding two equivalent subsets, as large as possible. While such an approach is more computationally taxing, it avoids some of the problems with the normalization or decomposition algorithms. This approach is taken in RTG Tools, described in a pre-print of Cleary *et al.* (2015). Their tool implements an exponential time exact algorithm, but uses clever bounding strategies to prune the search space and make the run-time feasible. To avoid blow-ups in runtime, it employs a cutoff strategy when the search space is too large to skip the matching of some variants. However, RTG Tools suffers from large RAM usage and can still fail to match variants in very dense regions, when the cut-off is activated.

Other related work includes Krawitz *et al.* (2010) and Assmus *et al.* (2013), which gives methods to check if two indels are equivalent, however, their analysis does not extend to matching non-singleton entry sets. Mäkinen and Rahkola (2013) and Mäkinen and Valenzuela (2014) describe a global approach for comparing two variant sets: create two donor genomes by inserting the respective variant sets, and measure the edit distance between them. Their approach is notable because it uses edit distance as an optimization criteria, as opposed to the number of matched variants; however, it has not been applied to mammalian sized genomes. Wittler *et al.* (2015) also studied the problem of matching variants that are large deletions (≥ 20bp), which is complementary to our study of small variants of different types.

In this paper, we present a divide-and-conquer algorithm for the variant matching problem. It is based on a theoretical result which shows how to partition the set of variants into small clusters which can be matched independently and in parallel. The partitioning step is linear in the number of variants, and, while the run-time is still exponential within each cluster, the size of each cluster is small in practice. VarMatch is more robust to different representation of complex variants and is able to detect more matches then the normalization or decomposition algorithms. It is also faster and uses an order-of-magnitude less memory then RTG Tools. We show that it is particularly useful in low complexity regions or those dense in variants.

Our theoretical result also allow VarMatch to seamlessly support different optimization criteria. VarMatch can maximize the number of variants matched, but can also maximize the sum of matched edit distances, which can increase robustness to different variant representations. Additional scoring schemes can also be easily integrated as long as there is a brute force algorithm for computing them—since partitioning divides the problem into small instances, the asymptotic running times of computing scores are rarely a factor. VarMatch can also support matching of VCF files that include genotype information to one that does not distinguish between hetero- and homozygous calls. Finally, the VarMatch software provides summary statistics, annotations, and visualizations that are useful for understanding callers’ performance.

## 2 DEFINITIONS

Let *x* be a sequence of elements (possibly a string). We use *x*[*i*] or *x_i_* to denote the element at position *i*, for 0 ≤ *i* < |*x*|, and we use *x*[*i, j*] to denote the sub-sequence *x_i_*, …*x_j_*, for 0 ≤ *i* ≤ *j* ≤ |*x*|. For two sequences *x* and *y*, we use the notation *x* · *y* or just *xy* to be the sequence obtained by their concatenation. A *tandem repeat* is a string (*x*_1_*x*_2_)^*m*^*x*_1_, where *x*_1_ and *x*_2_ are strings, *x*_2_ is non-empty, and *m* > 1 is an integer. We refer to *x*_1_*x*_2_ as the *repeat unit*.

Let *R* be a string, which we call the reference genome. A *variant* is a triple (*p, r*, **a**), where *p* is an index into *R, r* is a string and **a** is a pair of strings *a*_0_ and *a*_1_, with *r, a*_0_, *a*_1_ possibly empty. We refer to *r* as the reference allele and to **a** as the alternate alleles, and require that *r* = *R*[*p, p* + |*r*| − 1].

We exclude the possibility that both alternate alleles are the same as the reference allele, as this indicates no variation. However, it is possible that one of the alternate alleles is the same as the reference allele. It is also possible that neither of the alternate alleles are the same as the reference allele, meaning that the reference allele is not present in the donor. We refer to a variant with *a_0_* = *a_1_* as *homozygous* and as *heterozygous* otherwise – note that this is irrespective of the reference allele. In this paper, we assume the genome is diploid, but note that a haploid genome can be represented in our framework by setting *a_1_* = *a*_0_ for all variants.

We say that a variant *affects* the substring *R*[*p, p* + |*r*| − 1] of the reference genome. A sequence of variants *V affects* a given region of the reference if *V* contains at least one variant that affects that region. Let *v* = (*p, r*, **a**) and *v*′ = (*p*′, *r*′, **a**′) be two variants and assume without loss of generality that *p* ≤ *p*′. We say that *v* and *v*′ are *independent* if the intervals [*p, p* + |*r*|) and [*p*′, *p*′ + |*r*′|) do not overlap. In other words, *v* and *v*′ affect different regions of the reference genome.

We can *apply* a variant *v* = (*p, r*, **a**) to obtain two donor strings, as follows. There are three possible ways to incorporate a variant into the donor: it can be excluded or it can be included using one of two different ordering of the alleles. Let *c* ∈ {−1, 0, 1} be a *selection* value and let *j* ∈ {0, 1}. We define the selection function *s*(*v, c, j*) as *s*(*v*, 0, *j*) = *r, s*(*v*, 1, *j*) = *a_j_*, and *s*(*v*, −1, *j*) = *a*_1–*j*_. Applying *v* using *c* then gives two strings *d*_0_ and *d*_1_, where *d_j_* is obtained by replacing *R*[*p, p* + |*r*| − 1] with *s*(*v, c, j*).

Let *V* be a sequence of variants, and let Φ*_V_* = {0, 1, −1}^|*V*|^ be a *selection sequence*. A selection sequence Φ*_V_* is used to represent the selection for each variant in *V*. Suppose that for all *i* ≠ *k*, if |Φ*_V_* [*i*]| = |Φ*_V_* [*k*]| = 1, then *V*[*i*] and *V*[*k*] are independent. Then, we can *apply* the sequence of variants *V* using Φ*_V_* as follows. Let *D*(*R, V*, Φ*_V_*, *j*) represent the string obtained by applying all the variants simultaneously, where the *i*^th^ variant is applied using the selection function *s*(*V* [*i*], Φ*_V_* [*i*], *j*). Because of the independence condition, the final donor sequence obtained is the same regardless of what order the variants are applied in. Applying *V* using Φ*_V_* then gives us two strings: *D*(*R, V*, Φ*_V_*, 0) and *D*(*R, V*, Φ*_V_*, 1).

Consider two sequences of variants, *V* and *W*, and their corresponding selection sequences Φ*_V_*, Φ*_W_*. We say that (*V*, Φ*_V_*) and (*W*, Φ*_W_*) are *genotype equivalent* if *D*(*R, V*, Φ*_V_, j*) = *D*(*R, W*, Φ*_W_, j*) for all *j* ∈ {0,1}. On the other hand, some variant callers do not output genotype information. In these cases, a VCF entry indicates that the donor must contain at least one of the alternate alleles. For handling these types of datasets, we introduce the notion of variant equivalence. Variant equivalence is similar to genotype equivalence, but two variants are considered matched if they share at least one alternate allele. Formally, (*V*, Φ*_V_*) and (*W*, Φ*_W_*) are *variant equivalent* if *D*(*R, V*, Φ*_V_*, 0) = *D*(*R, W*, Φ*_W_*, 0).

Given a variant *v* = (*p, r*, **a**) and a selection value *c*, the score of *v* in the *unit cost model* is |*c*|. In the *edit distance cost model*, its score is |*c*|(*E*(*r, a*_0_) + *E*(*r, a*_1_)), where *E*(·) is the edit distance function. For a variant sequence and a selection sequence, the score is the sum of the scores for each variant. Suppose we have two variant sequences *V,W* and their selection sequences Φ*_V_*, Φ*_W_*. Under the *baseline scoring scheme*, their score is the score of *W*, while in the *total scoring scheme*, their score is the score of *V* plus the score of *W*. Note that we can compute the score even if (*V*,Φ*_V_*) and (*W*,Φ*_W_*) are not equivalent.

In sum, we can define four different scoring functions *F* (*V,W*,Φ*_V_*,Φ*_W_*). These correspond to a choice of the unit vs. edit distance cost model and the baseline vs. total scoring schemes. The baseline scoring scheme is appropriate when comparing multiple datasets against one ground truth, while the total scoring scheme is more appropriate for a two-way comparison of different tools when a ground truth is not available. The unit distance cost model is traditionally used, but, unlike the edit distance cost model, is not robust to the decomposition of complex variants. For example, a single complex variant counts the same as an equivalent entry set of three SNVs (Figure 1(b)). We believe that matching using both models can make any resulting conclusions more robust to diversity in variant representation and/or highlight important differences.

Given a reference genome *R*, two variant sequences *V* and *W*, a type of equivalence (either genotype or variant), and a scoring function *F*, the variant matching problem VarMatch (*R, V, W*) is to find two corresponding selection sequences (Φ*_V_*, Φ*_W_*) such that (*V*, Φ*_V_*) and (*W*, Φ*_W_*) are equivalent and have the highest score amongst all equivalent pairs. Intuitively, in the unit cost model, maximizing the score results in trying to match as many variants as possible. In the edit distance cost model, maximizing the score results in trying to match as many nucleotide differences with the reference genome as possible.

## 3 METHODS

In this section, we develop an algorithm for the VarMatch problem. First, we prove Theorem 1, which allows a divide-and-conquer strategy to be applied (Section 3.1). Then, we describe our algorithm to partition a problem into smaller sub-problems using the LinearClustering algorithm (Section 3.2). Finally, we solve each subproblem using an exact branch and bound algorithm, based on Cleary *et al.* (2015). This is done over multiple threads, with each thread solving its own subproblem. The computational complexity of VarMatch remains open.

### 3.1 Partitioning theorem

In this section, we derive our main theorem which forms the basis of the VarMatch algorithm. First, we prove the following two lemmas about strings.

#### Lemma 1.

*Given non-empty strings a, b, and c, if ab* = *bc, then d* = *abc is a tandem repeat with repeat unit a*.

#### Proof.

We prove by induction on the length of *d*. The base case of |*d*| = 3 is trivial. In the general case, consider three possibilities. In the case that | *b*| = |*a*|, then *a* = *b* = *c* and *d* = *a*^3^ is a tandem repeat with *a* is a repeat unit.

Now consider the case that |*b*| < | *a*|. Then, *b* is a prefix of *a*, and we can write *a* = *bb*_0_, for some non-empty string *b*_0_. Then, *bc* = *ab* = *bb*_0_*b*, and *c* = *b*_0_*b*. Then *d* = *abc* = *bb*_0_*bb*_0_*b* is a tandem repeat with repeat unit *bb*_0_ = *a*.

Now consider the case that |*b*| > |*a*|. Since *ab* = *bc*, then |*a*| = |*c*|. Moreover, *b*’s prefix of length |*b*| − |*c*| is equal to its suffix of length |*b*| − |*a*|. Denote this string as *b*_1_. We can then write *b* = *ab*_1_ and *b* = *b*_1_ *c*. We now apply the Lemma inductively to the equality of *ab*_1_ = *b*_1_ *c* and get that *ab*_1_ *c* is a tandem repeat with repeat unit *a*. Then, *d* = *abc* = *aab*_1_ *c* is also a tandem repeat with repeat unit *a*.

#### Lemma 2.

*Given strings s*_1_ = *x*_1_*yz*_1_ *and s*_2_ = *x*_2_*yz*_2_, *if x*_1_ ≠ *x*_2_ *or z*_1_ ≠ *z*_2_, *y is not a tandem repeat and* |*y*| > 2 · min(*abs*(|*x*_1_| − |*x*_2_|), *abs*(|*z*_1_| − |*z*_2_|)), *then s*_1_ ≠ *s*_2_.

#### Proof.

First, observe that if either |*x*_1_| = |*x*_2_| or |*z*_1_| = |*z*_2_|, then it is trivial to show that *s*_1_ ≠ *s*_2_. Therefore, we can assume that |*x*_1_| = |*x*_2_| and |*z*_1_| = |*z*_2_|. Also assume, without loss of generality, that |*x*_1_| > |*x*_2_|.

Assume for the sake of contradiction that *s*_1_ = *s*_2_. Then, since |*s*_1_| = |*s*_2_|, we have that |*x*_1_| − |*x*_2_| = |*z*_2_| − |*z*_1_|. Therefore, |*y*| > 2 · min(*abs*(|*x*_1_| − |*x*_2_|), *abs*(|*z*_1_| − |*z*_2_|)) = 2(|*x*_1_| − |*x*_2_|). We can then divide *y* into three non-empty parts *y* = *y*_1_*y*_2_*y*_3_, such that |*y*_1_| = |*y*_3_| = |*x*_1_| − |*x*_2_|. Because *s*_1_ = *s*_2_, we have that *x*_1_*y*_1_*y*_2_*y*_3_*z*_1_ = *x*_2_*y*_1_*y*_2_*y*_3_*z*_2_. Using what we know about the lengths of these strings, we have that *y*_1_*y*_2_ = *s*_1_[|*x*_1_|, |*s*_1_ — |*z*_1_| − |*y*_3_| − 1]] = *s*_1_ [|*x*_1_|, |*s*_1_| − |*z*_2_| − 1]] = *s*_2_[|*x*_1_| |*s*_2_| |*z*_2_| − 1]] = *s*_2_[|*x*_2_| + |*y*_1_|, |*s*_2_| − |*z*_2_| − 1]] = *y*_2_*y*_3_. Applying Lemma 1 to the equality *y*_1_*y*_2_ = *y*_2_*y*_3_, we get that *y* = *y*_1_*y*_2_*y*_3_ must be a tandem repeat, contradicting the conditions on *y*.

Let *X* and *Y* be two variant sequences. We define the *max change in length* of *X* and *Y*, denoted by MCL(*X, Y*), as

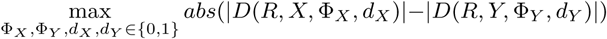

Intuitively, MCL(*X, Y*) is the maximum difference in the size of the donor sequences that can be obtained for any selection sequences over *X* and *Y*.

#### Theorem 1.

*Consider an interval of the reference*, [*b, e*], *for* 0 ≤ *b* ≤ *e* < |*R*|. *Consider four variant sequences, V_i,j_, for i* ∈ {0, 1} *and j* ∈ {0, 1}, *satisfying*

1. *V*_0,0_ *and V*_1,0_ *only affect R*_0_ = *R*[0, *b* − 1],
2. *V*_0,1_ *and V*_1,1_ *only affect R*_1_ = *R*[*e* + 1, |*R*| − 1].
3. *R*[*b, e*] *is not a tandem repeat*
4. |*R*[*b, e*]| > 2 · min*_j_* (*MCL*(*V*_0,*j*,_ *V*_1,*j*_))

*For all j, let* (Φ_0,*j*_, Φ_1,*j*_) *be an optimal solution for* VarMatch(*R_j_, V*_0,*j*_, *V*_1,*j*_). Let *V_i_* = *V*_*i*,0_ · *V*_*i*,1_ *and Φ_i_* = Φ_*i*,0_ · Φ_*i*,1_, *for all i. Then*, (Φ_0_, Φ_1_) *is an optimal solution for* VarMatch (*R, V*_0_, *V*_1_).

#### Proof.

Let 
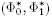
 be an optimal solution for VarMatch(*R, V*_0_, *V*_1_), and suppose for the sake of contradition that it is better than (Φ_0_, Φ_1_).

Let 
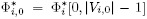
 and 
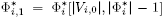
. For the score functions we consider in this paper, 
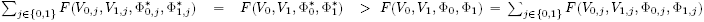
. Then, there must exist a *j* such that 
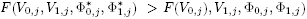
. We can assume without loss of generality that *j* = 0.

Intuitively, this means that the optimal solution to VarMatch(*R, V*_0_, *V*_1_), projected onto the “left” variants (i.e. *V*_0,0_ and *V*_1,0_), has a higher score then the optimal solutions of the “left” problem alone, i.e. VarMatch(*R*_0_, *V*_0,0_, *V*_1,0_). This can only be possible because this projection is not a feasible solution to the “left” problem, i.e. the donor sequences are not identical. Thus, there exist *d* ∈ {0,1} such that 
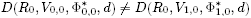
.

Let 
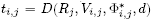
 and 
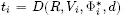
, for all *i* and *j*, We will apply Lemma 2 to the strings *t*_1_ and *t*_2_. To see that the conditions of Lemma 2 apply, observe that

- ∀*i, t_i_* = *t*_*i*,0_ · *R*[*b, e*] · *t*_*i*,1_
- *t*_0,0_ ≠ *t*_1,0_
- *R*[*b, e*] is not a tandem repeat.
- |*R*[*b, e*]| > 2·min*_i_*(MCL(*V*_0,*j*_, *V*_1,*j*_)) ≥ 2·min*_j_, abs*(|*t*_0,*j*_| − |*t*_1,*j*_|)

Then, Lemma 2 implies that *t*_0_ ≠ *t*_1_. However, this contradicts that 
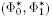
 is a solution for VarMatch(*R, V*_0_, *V*_1_).

The following counterexample shows that Theorem 1 is not true if *R*[*b, e*] is a tandem repeat. Let *R* = CCATATATGC be the reference sequence, let *b* = 2 and *e* = 7, and let

- *V*_0,0_ = (0, CC, ∊) and *V*_1,0_ = (0, CC, AT)
- *V*_0,1_ = (7, GC, AT) and *V*_1,1_ = (7, GC, ∊)

Here, ∊ denotes the empty string. Observe that all conditions of the Theorem are satisfied, except that *R*[*b, e*] = ATATAT is a tandem repeat. In particular, |*R*[*b, e*]| = 6 > 2 · min*_j_*(MCL(*V*_0,*j*_, *V*_1,*j*_)) = 4. The optimal solutions for the two corresponding VarMatch subproblems do not select any variants, and the Theorem implies that the optimal solution to VarMatch(*R, V*_0_, *V*_1_) is also empty. However, the optimal solution selects all variants since {*V*_0,0_, *V*_0,1_} and {*V*_1,0_, *V*_1,1_} are equivalent.

### 3.2 Clustering algorithm

Theorem 1 can be applied to the VarMatch problem to divide the input into subproblems that can be solved separately. In particular, we can identify *separator* regions — long-enough regions of the reference that are not affected by any variants and are not tandem repeats. We can then divide our variants into two clusters — those on the left and on the right of the separator region. Each cluster can be solved independently, and the solutions can then be trivially combined.

Many clustering strategies are possible, based on how these separator regions are identified. We found a simple greedy strategy works well in practice. We make a linear scan through all the variants, and for each new variant, we check if the reference region between it and the previous variant is a separator. If it is, then we start a new cluster and add the current variant to it. If not, we simply add the current variant to the current cluster. The pseudocode for the algorithm, called LinearClustering, is provided below.

**Algorithm.**
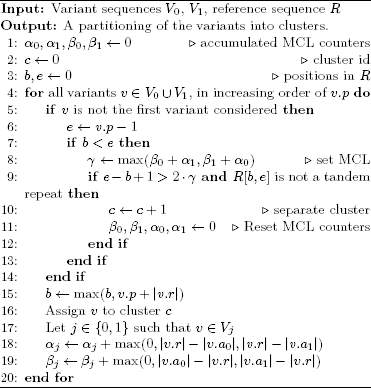
Linear Clustering

In order to check if a region is long-enough to be a separator, we check if its length is > 2*γ*, where 
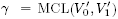
 and 
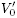
 and 
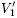
 are the variants in the current cluster. This ignores the MCL of the variants to the right of the separator, which are not yet known due to the greedy nature of our algorithm. In terms of Theorem 1, instead of finding min*_j_* MCL(*V*_0,*j*_, *V*_1,*j*_), our algorithm just calculates MCL(*V*_0,0_, *V*_1,0_). Theorem 1 still applies, since |*R*[*b, e*]| > 2 · MCL(*V*_0,0_, *V*_1,0_) ≥ min*_j_* MCL(*V*_0,*j*_, *V*_1,*j*_).

To maintain the current value of *γ*, our algorithm maintains four counters. The running total of the maximum possible decrease (respectively, increase) of the reference length for the variants 
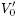
 and 
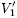
 is maintained by *α*_0_ and *α*_1_ (respectively, *β*_0_ and *β*_1_). Then, we can compute *γ* = max(*β*_0_ + *α*_1_, β_1_ + *α*_0_). To check if a potential separator is tandem repeat, we use a simple algorithm described in Fungtammasan *et al.* (2015).

Finally, we solve each subproblem using a variation of the exact branch and bound algorithm of Cleary *et al.* (2015). Due to its high similarity, we do not describe it here, but, for the sake of completeness and clarity, we provide its details in the Supplementary Text. The algorithm’s running time and memory usage is Ω(3^|*V*| + |*W*|^). However, the algorithm is fast in practice, since it is applied only on small subproblems generated by the LinearClustering algorithm and it employs pruning strategies.

## 4 RESULTS

We implemented the VarMatch Toolkit, which takes multiple query VCF files and matches them separately to one baseline VCF file. The baseline could be a ground truth set, or, if one is unavailable, any of the callsets. Based on the various scoring functions and equivalence definitions given in Section 2, our software runs in different modes. The cost model of VarMatch can be either unit (denoted by U) or edit distance (E). The equivalence mode can be either genotype (G) or variant (V). Also, the scoring scheme can be either baseline (B), query (Q), or the total (T). The query scoring scheme is applicable when there is only one query file and seeks to maximize the score of the query. It is like the baseline scoring scheme defined in Section 2 but with the roles of query and baseline reversed. Considering all combinations of above, there are 12 possible modes, each denoted by a three letter abbreviation (e.g. UGT for unit cost, genotype equivalence, and total scoring scheme).

For each query, VarMatch automatically matches it to the baseline, simultaneously using all the modes. VarMatch then outputs files containing annotations of matched variants and recall and precision statistics and plots. It also identifies and outputs visualizations of variants that are matched in one mode but not in another (as in Supplementary Figure 2). VarMatch can also create Precision-Recall curves by varying the minimum quality cutoff for the VCF file (not shown).

### 4.1 Datasets

To evaluate the performance of VarMatch, we selected five state-of-the-art variant calling methods: Platypus(pt), GATK HaplotypeCaller(hc), GATK UnifiedGenotyper(ug), Freebayes(fb) and SAMtools(st). The algorithms of these tools vary, resulting in potentially different representations of variants. For example, pt performs local assembly, pt/hc/ug perform local realignment, and pt/fb phase haplotypes and thus tend to merge variants into longer complex variants.

For evaluation, we use the variant call sets provided by Li (2014). The CHM1 dataset comes from 65x Illumina sequencing of the haploid CHM1hTERT cell line, mapped to GRCh37 using bowtie2, and small variants called separately by fb/hc. The CHM1 dataset has less variants, due to the haploid nature of the CHM1hTERT cell line. The NA12878 dataset comes from 55x Illumina sequencing of the NA12878 diploid cell line, mapped to GRCh37 using bowtie2, and small variants called separately by fb/ug/pt/st. It also includes Freebayes run on mappings generated by BWA-MEM, which we will refer to as bwa-fb. Samtools was not run in genotype mode, and only the presence of alternate alleles, and not the corresponding genotype, was reported.

### 4.2 Evaluating VarMatch’s accuracy and resources

We compared VarMatch against the normalization algorithm (running ‘normalize’ function of vt(version v0.5) (Tan *et al.*, 2015) followed by strict matching), the decomposition algorithm (running ‘vcfallelicprimitives’ function of vcflib, followed by normalization algorithm and strict matching) and RTG Tools (Cleary *et al.*, 2015) (version 3.6.2, running ‘vcfeval’ function with paramaters ‘-all-records’ and ‘-ref-overlap’). The decomposition algorithm changes the number of variants in a VCF file and thus poses a challenge for counting the number of matches. In this case, an original query entry (the baseline case is symmetric) is said to be *partially matched* if at least one of its decomposed child variants is matched but either 1) one of its decomposed children is not matched, or 2) one of its decomposed children matches a baseline decomposed child which has a decomposed non-matched sibling. Partial matches do not reflect true matches and thus are not counted. All experiments were run on an Intel Xeon CPU with 512 GB of RAM and using 8 cores (at 2.67GHz). VarMatch and RTG Tools are multi-threaded and were allowed to use all threads, while vt and vcflib are single threaded. VarMatch was run in UGT mode unless otherwise stated.

Tables 1 and 2 show the exact matching results of comparing CHM1 fb to CHM1 hc datasets, and the NA12878 fb and ug datasets. Note that since we only consider exact matches, the algorithm that can find the highest number of matches is the best. The variants matched by the decomposition algorithm were always a subset of the variants matched by RTG Tools, which were in turn always a subset of the variants matched by VarMatch. RTG Tools and VarMatch match the same set of variants in Table 1, but in Table 2, RTG Tools reported skipping 23 genome regions because it reached the search space upper bound, discarding all the variants in these regions. As a result, VarMatch matches more entries then RTG Tools. Though the number of these variants is small, they are located in regions dense with variants which might be of particular interests to researchers. Both RTG Tools and VarMatch match more entries than the normalization or decomposition algorithms, but VarMatch uses less running time and an order of magnitude less memory than RTG Tools.

**Table 1.**
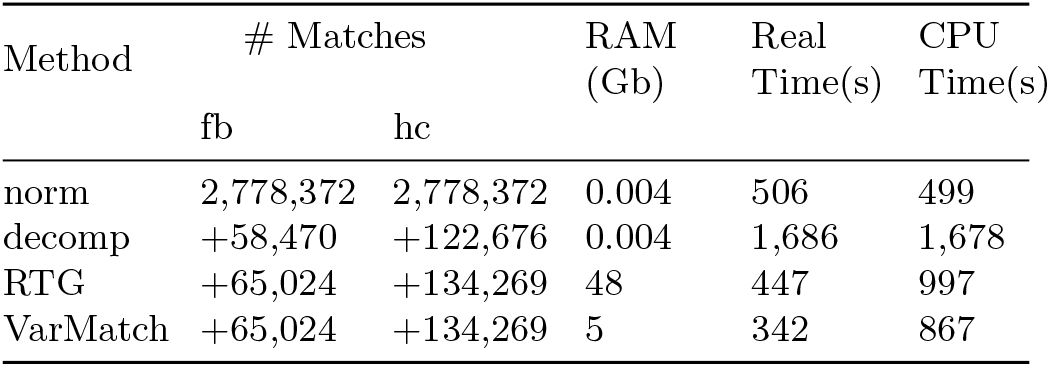
Comparison of VarMatch to CHM1 fb and hc datasets. Numbers of matches are shown as offsets to the baseline numbers of the normalization algorithm. We measure both the real running time and the total number of CPU-seconds that the process spent. Normalization and decomposition are single threaded, so the real time matches closely the CPU time.

| Method | # Matches |  | RAM<br>(Gb) | Real<br>Time(s) | CPU<br>Time(s) |
| --- | --- | --- | --- | --- | --- |
|  | fb | hc |  |  |  |
| norm | 2,778,372 | 2,778,372 | 0.004 | 506 | 499 |
| decomp | +58,470 | +122,676 | 0.004 | 1,686 | 1,678 |
| RTG | +65,024 | +134,269 | 48 | 447 | 997 |
| VarMatch | +65,024 | +134,269 | 5 | 342 | 867 |

**Table 2.**
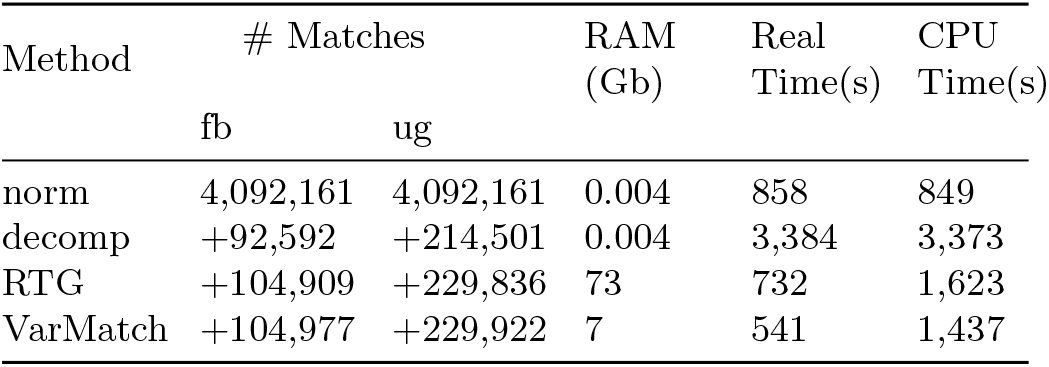
Comparison of VarMatch to NA12878 fb and ug.

We also evaluated the effectiveness of our clustering algorithm. Figure 2 shows the distribution of the sizes of each subproblem. VarMatch partitions 6,438,208 initial small variants into 3,272,206 subproblems, 99.9% of which have less than nine variants in them.

**Fig. 2:**
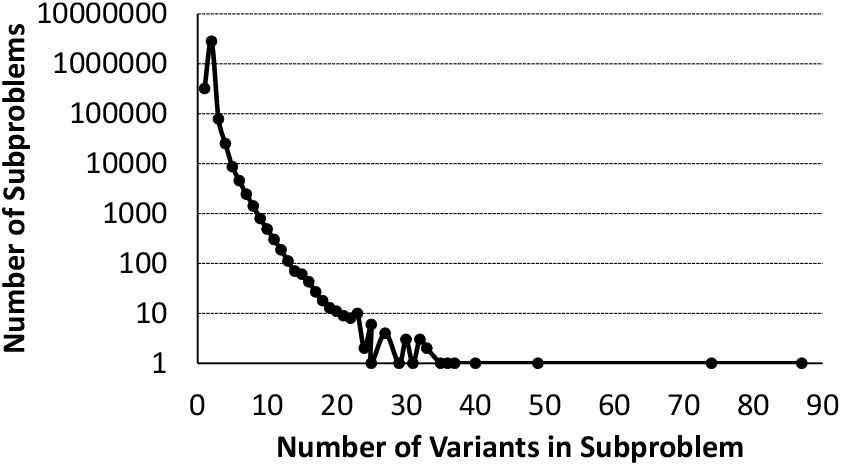
Distribution of number of variants in subproblems for experiment in Table 1.

### 4.3 Comparison of different modes

To test VarMatch’s ability to match between datasets with and without genotype information, we use UVT mode to compare the NA12878 fb and st datasets against the GIAB benchmark (version 2.18) (Zook *et al.*, 2014). We find that fb has a precision of 49.85% and a recall of 99.64%, while st has a precision of 63.90% and a recall of 98.96%. RTG Tools could not process the st dataset due to lack of genotype calls.

We then evaluate the difference in results when different scoring schemes and equivalence modes are used. Table 3 illustrates the results on the NA12878 fb and ug datasets. Observe that on this dataset, ug outperforms fb in most modes, but there is a discrepancy when variant equivalence is used — the edit distance score favors fb while the number of variants matched favors ug. This could indicate that, in this dataset, fb is better at detecting the presence of variants then it is at genotyping them. Observe also that fb has more matches when the scoring scheme maximizes the number of fb matches (e.g. UGB) then when the total number of matches in both fb and ug are maximized (e.g. UGT). VarMatch flags variants that are matched in one mode but not in the other, making it possible for a researcher to further investigate the source of such discrepancies. These may uncover bugs, quirks, or features of a variant caller. Supplementary Figures 1 and 2 show examples, simplified from real data, that illustrate why the number of matched variants varies under different criteria. We recommend users to run VarMatch simultaneously in all modes (the default option), and using the graphs, tables, and visualizations provided in the output to detect and understand any anomalies in the datasets.

**Table 3.** Number of matched variants and the edit distance score from the NA12878 fb and ug datasets, under different modes of VarMatch. Baseline is arbitrarily chosen to be fb, and the query to be ug. The top half shows results in genotype equivalence modes and the bottom half in variant equivalence modes. Numbers are given as offsets to UGT in top half and to UVT in bottom half.

| Mode | # Matched Entries |  | Edit Distance Score |  |
| --- | --- | --- | --- | --- |
|  | fb | ug | fb | ug |
| UGT | 4,197,138 | 4,322,083 | 5,184,864 | 5,169,151 |
| UGB | +8 | -32 | +31 | -12 |
| UGQ | -21 | +6 | -115 | -36 |
| EGT | +8 | -15 | +35 | +15 |
| EGB | +8 | -45 | +43 | -22 |
| EGQ | -6 | -9 | +15 | +18 |
| UVT | 4,266,505 | 4,411,621 | 7,042,027 | 7,046,191 |
| UVB | +33 | -913 | -942 | -1,646 |
| UVQ | -84 | +15 | -166 | -30 |
| EVT | -8 | -775 | +85248 | +85280 |
| EVB | -4 | -998 | +85382 | +84366 |
| EVQ | -70 | -740 | +84844 | +85346 |

Finally, we evaluated the robustness of matching results to the score model being used. We repeated several unit cost model experiments from the paper in the edit distance cost model (Supplementary Table 1). We found that there was little difference in the results.

### 4.4 Matching variants in hard regions

To illustrate the power of VarMatch to detect matches, we focus on genomic regions where variants are particularly hard to match.

We downloaded co-ordinates of low complexity regions (Li, 2014), covering about 2% of the autosomal genome. Alignment and variant calling is particularly challenging in these regions, often leading to different representations, and we evaluated how VarMatch performs there. Table 4 shows the comparison of NA12878 ug, pt, and bwa-fb datasets, restricted to variants in the low complexity regions. Because no ground truth is available in these regions, we arbitrarily used ug as a baseline. The goal is to identify all the high-confidence variants, i.e. thost that occur in all three datasets. VarMatch is able to detect 14% more high-confidence variants than the normalization algorithm, and 4.8% more than the decomposition algorithm, and 576 more (.16%) than RTG Tools. RTG Tools skipped some genome regions because of its search space upper bound and was also unable to process some of the genotype information from pt.

**Table 4.**
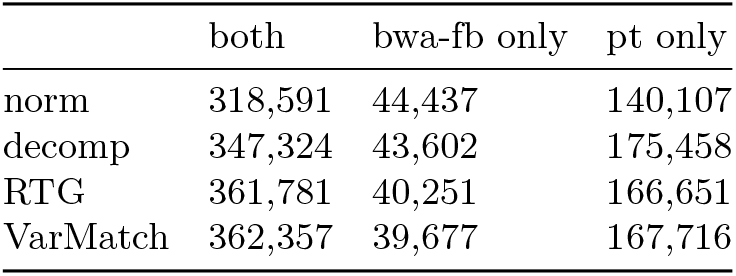
Number of baseline variants matched in low complexity regions. VarMatch was run in UGB mode using ug as the baseline. For each variant matching experiment, baseline variants are categorized as being matched in both bwa-fb and pt datasets or in just one of the two datasets.

|  | both | bwa-fb only | pt only |
| --- | --- | --- | --- |
| norm | 318,591 | 44,437 | 140,107 |
| decomp | 347,324 | 43,602 | 175,458 |
| RTG | 361,781 | 40,251 | 166,651 |
| VarMatch | 362,357 | 39,677 | 167,716 |

A recent evaluation study of Highnam *et al.* (2015) compared the results of different variant callers and aligners against the GIAB benchmark (Zook *et al.*, 2014), However, they excluded dense regions – any 10bp regions that contain an indel and another variant in the benchmark – due to the difficulty of matching variants in those regions. However, the accurate detection of variants in such dense regions is particularly important in studying the mechanisms that give rise to variation. For example, the presence of a cluster of small events near structural variation breakpoints can help differentiate microhomology-mediated break-induced replication from non-homologous end joining (Hastings *et al.*, 2009; Mäkinen and Rahkola, 2013). Meanwhile, some variants in dense regions are disease related, e.g. 51 variants from dense regions in the HLA region.

In Table 5, we measured the accuracy of fb and pt in detecting specifically the variants in dense regions. On the whole benchmark, fb has higher recall than pt, but pt has higher recall in dense regions, using VarMatch (Table 5) and RTG Tools (Supplementary Table 2). The use of VarMatch or RTG Tools can therefore allow studies such as Highnam *et al.* (2015) to measure accuracy in these important regions.

**Table 5.** The number and recall of benchmark variants matched by fb and pt (in UGB mode), from the whole genome (first row) and just the dense regions (second row).

|  | # Matched Benchmark Entries (Recall) |  |
| --- | --- | --- |
|  | fb | pt |
| genome-wide | 2,896,841 (99.35%) | 2,891,849 (99.18%) |
| dense regions | 24,188 (84.69%) | 24,522 (85.86%) |

## 5 DISCUSSION

In this paper, we presented VarMatch, an open-source parallel tool for matching equivalent genetic variants. VarMatch is robust to different representations of complex variants and supports flexible scoring schemes. We demonstrated that it can detect more matches than the normalization algorithm and is faster and uses an order-of-magnitude less memory than RTG Tools. It is important to note that, for evaluating a variant caller, VarMatch should be used in conjunction with other validations (e.g. longer haplotypes are not reflected in a higher match score but are a desirable feature).

There are efforts underway to represent the reference as a graph instead of as a string, led by the GA4GH coaliation. This would affect variant matching algorithms, which would need to adopt to new reference formats; however, such formats are not yet stabilized. For the time being, most studies continue to use a string as a reference and require accurate variant matching tools.

One weakness of VarMatch is its worst-case exponential running time. There are theoretical cases when the runtime would become infeasible, when there is large number of similar variants within a long tandem repeat region (e.g. 100 SNVs within a 500bp poly-A region). In such a case, our clustering algorithm would fail to divide the input and the branch and bound algorithm would also fail to prune the search space. We did not observe such extreme conditions in our experiments, however, to handle this contingency, we monitor our search space and when it reaches a fixed upper bound, we apply the strict matching algorithm to the cluster. For the same reason, VarMatch is limited in processing population-scale call sets like dbSNP, where variants are densely packed and our clusters become large. In such situations, the decomposition algorithm can be used.

Finally, the power of Theorem 1 is not fully explored in our linear clustering algorithm. For instance, it might be possible that while the region bounded by two variants is not a separator, a smaller sub-region is. Additionally, we only calculate MCL(*V*_0,0_, *V*_1,*j*_), but computing minj MCL(*V*_0,*j*_, *V*_1,*j*_) may detect new separators. Our current linear clustering algorithm is sufficient for our experiments, but a more powerful clustering algorithm is also theoretically possible.

## Acknowledgements

This work has been supported in part by NSF awards DBI-1356529, CCF-1439057, IIS-1453527, and IIS-1421908 to PM.

